# Filamentous growth, cell envelope architecture, and surface appendages of a member of the Chloroflexota, *Litorilinea aerophila*

**DOI:** 10.1101/2025.08.19.671072

**Authors:** Marie Joest, Clara L. Mollat, Laura Mutschler, Marta Rodriguez-Franco, Shamphavi Sivabalasarma, Friedel Drepper, Pitter F. Huesgen, Thomas Ott, Sonja-Verena Albers

## Abstract

*Litorilinea aerophila*, a filamentous bacterium of the phylum Chloroflexota (class *Caldilineae*), exhibits unique morphological and cell envelope features that challenge traditional bacterial models. Initially described as Gram-negative, Chloroflexota are increasingly considered as monoderm, lacking a true outer membrane. In this study, we investigated the growth behaviour, cell morphology, and cell appendages of *Litorilinea aerophila* using light microscopy, cryo-electron tomography, and structural biology. Dry weight-based growth assays revealed a prolonged lag phase (~40 h) followed by exponential and stationary phases. Light and fluorescence microscopy revealed irregular indentations along cell filaments, accompanied by a diffuse distribution of DNA, indicating a multicellular organization. Thin-section electron microscopy confirmed septa formation, and in late growth stages, filaments became shorter with more defined indentations and membrane vesicle release. Next to the already characterised bacterial archaellum of *Litorilinea aerophila*, two additional types of surface appendages were identified: (i) pilus-like structures consistent with Tad pili, and (ii) grappling hook like structures. These findings contribute to the understanding of cell envelope diversity, growth, and surface structures in filamentous Chloroflexota.

## Introduction

Several members of the phylum of Chloroflexota, including representatives of the classes *Chloroflexi, Anaerolineaea, Caldilineae*, and *Ktedonobacteria*, form multicellular filaments and exhibit a distinctive, atypical cell envelope architecture (Sutcliffe, 2011; Hanada, 2014). The cell envelope of Chloroflexota has been of great interest as, despite gram-negative staining, they are considered monoderm. Despite the multilayered envelope architecture of some Chloroflexota, clear evidence for a lipid outer membrane is not available (van der Meer *et al*., 2002; Sutcliffe, 2011). The absence of genes of the beta-barrel assembly machinery (BAM-complex) and liposaccharide biosynthesis pathway, which are encoded in all known diderm phyla, supports the lack of an outer membrane (Sutcliffe, 2010). Instead, it is suggested that the layered structures are composed of polysaccharide or proteins (Sutcliffe, 2011). However, recent cryo-electron tomograms of *Candidatus* Viridilinea mediisalina and *Chloroflexus aggregans*, show a diderm-like envelope, yet they lack diderm signature pathways and genes. This suggests that Chloroflexota might harbor mono- and neoderm architecture that needs to be elucidated (Gaisin *et al*., 2020; Benyahia *et al*., 2025).

Archaea and Bacteria possess a diverse array of cell surface structures that mediate critical biological processes such as cell motility, DNA exchange, or biofilm formation. In bacteria, cell surface appendages are typically classified in two major categories: flagella and pili (Kim, 2017). Flagella are long, thick filamentous protein structures with a diameter of ~20 nm responsible for swimming motility (Beeby *et al*., 2020). In contrast, pili are thinner, hair-like structures that facilitate diverse functions including adhesion, cell-cell contact, and DNA exchange (Thanassi *et al*., 2012; Chaudhury *et al*., 2018).

Unlike bacteria, archaea do not use flagella for swimming motility. Instead, they encode a distinct motility structure called the archaellum. Despite serving a similar function to the bacterial flagellum, the archaellum is not evolutionarily related to it. Rather, the archaellum is a member of the type IV filament (TFF) superfamily—a diverse group of nanomachineries characterized by conserved core components and variable functional adaptations (Albers and Jarrell, 2018; Jarrell *et al*., 2022). The archaellum is homologous to the type IV pilus (T4P), a member of the TFF (Denise *et al*., 2019). T4P facilitate processes including motility, DNA uptake and adherence in bacteria and archaea. Another example for members of the TFF is the tight adherence (Tad) pilus, an archaea-derived T4P found in both bacteria and archaea. The tad gene cluster is widespread among archaea, likely due to horizontal gene transfer events (Kachlany *et al*., 2000; Denise *et al*., 2019). Although the archaellum was for a long time considered an archaea-specific motility machinery, recent discoveries have challenged this view. Several members of the bacterial phylum *Chloroflexota* have been found to harbor *bona fide* archaellum gene clusters, acquired through horizontal gene transfer (Hug *et al*., 2013; Lim *et al*., 2023; Sivabalasarma *et al*., 2025).

Motile Chloroflexota bacteria harbor flagella or archaella for swimming motility. Next to this, pilus-like structures were found in filamentous *C. aggregans* and *C. islandicus* (Fukushima *et al*., 2016; Gaisin *et al*., 2017). Recently, cryo-electron tomography also revealed pilus-like structures in *Roseiflexus castenholzii* and *Candidatus* Viridilinea mediisalina which are considered to be Tad pili as the genomes of *R. castenholzii* and *Candidatus* Viridilinea mediisalina only encode for genes for tad pilus biogenesis such as *rcpC-tadZABC* and no genes for other pilus biogenesis (Gaisin *et al*., 2020).

Beyond TFF systems, other classes of cell surface appendages have evolved to support unique ecological strategies. A striking example is found in filamentous *Aureispira* spp. such as *A. CCB-QB1*, which use grappling hook like structures to attach to the flagella of prey organisms (e.g. *Vibrio campbellii*). These filaments are unrelated to TFFs and are thought to be structurally linked to the type IX secretion system (T9SS). This predatory interaction is part of a specialized feeding strategy termed ixotrophy, in which the prey is immobilized and lysed via a Type VI secretion system. Ixotrophy provides *Aureispira* with a survival advantage under nutrient-limited conditions, highlighting the functional breadth of surface appendages in prokaryotic life (Lien *et al*., 2024).

Previously, we solved the first structure of a bacterial archaellum isolated from *Litorilinea aerophila*, a member of the class *Caldilineae* of the phylum Chloroflexota (Sivabalasarma *et al*., 2025). Here, we present a comprehensive characterization of *Litorilinea aerophila* focusing on growth characteristics, cell division, and cell surface filaments. We analyzed the growth of the filamentous bacteria over time and also cell division events. Additionally, we observed two more cell surface filaments, one pili-like structure and one grappling hook looking like structure, of which we solved the structure using cryoEM.

## Results

### Filamentous growth and morphology dynamics of *Litorilinea aerophila*

When cultivated in marine broth medium, *Litorilinea aerophila* exhibited a characteristic filamentous morphology previously described by Kale et al (2013). Due to extensive clump formation and the filamentous nature of the culture, standard optical density measurements were not possible for growth quantification. Instead, dry weight was measured over a period of 5 days (Figure 1A). To monitor morphological changes throughout growth, cells were analyzed at regular intervals via light and fluorescence microscopy. DNA and membrane dyes revealed irregularly spaced membrane indentations within the filaments, suggestive of a multicellular organization (Figure 1B). Diffuse DNA staining between membrane indentations further supported the interpretation of a multicellular organization, indicating the presence of compartmentalized nucleoids within individual filaments (Figure 1B, bottom row). Filament lengths frequently exceeded 100 µm.

**Figure 1:**
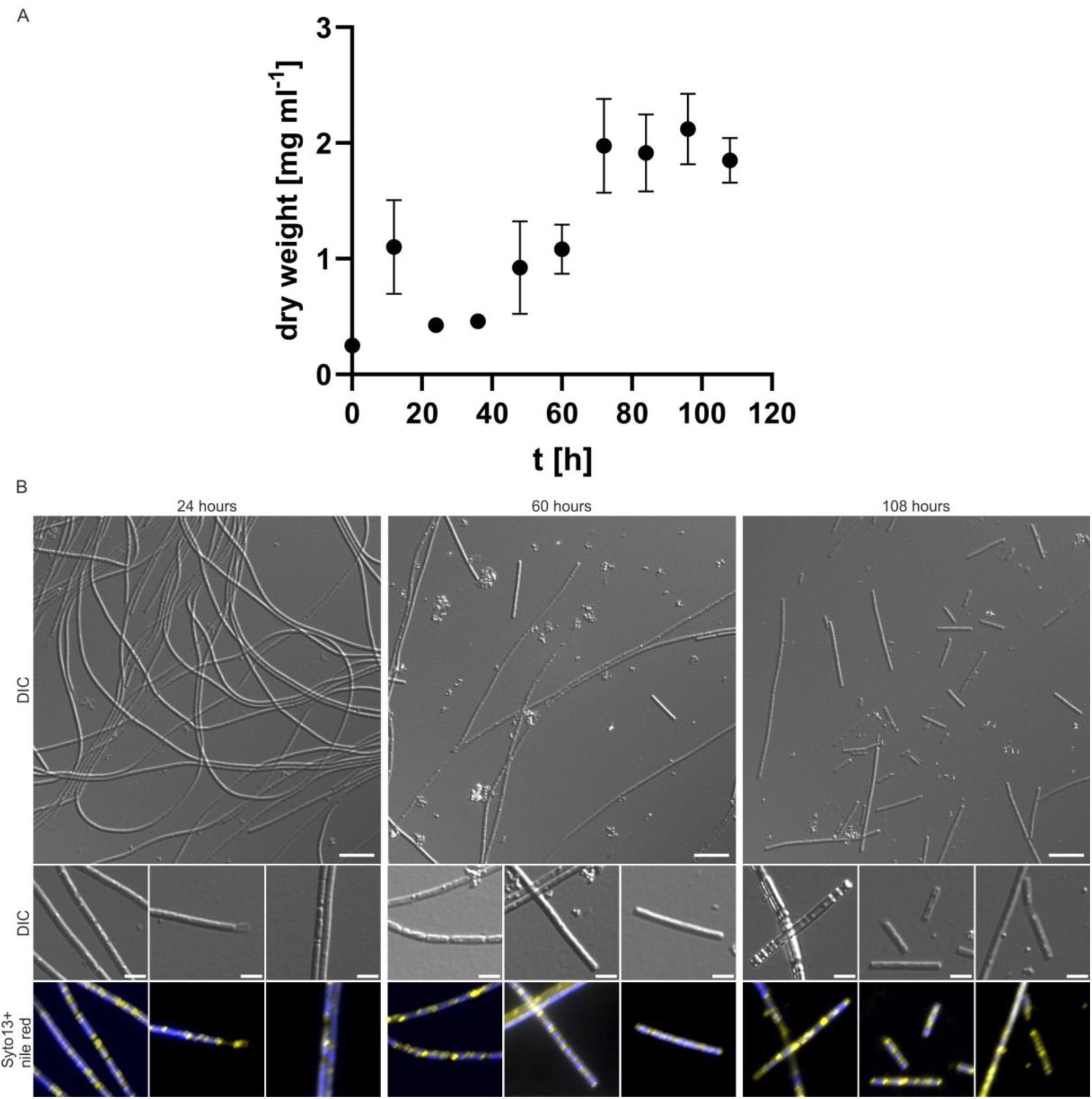
Cell growth and light microscopy images of *L. aerophila* cells cultured in marine broth medium. (A) Growth of *L. aerophila* measured in dry weight over 5 days. Each value represents the mean of three independent, biological replicates. Error bars show SEM. (B) Fluorescent light microscopy of L. aerophile cells after 24, 60 and 108 hours of growth. The membrane is stained with Nile red (shown in yellow), DNA is stained with Syto13 (shown in blue). Scale bars of the overview: 10 µm. Scale bar of the zoomed images: 3 µm.

Transmission electron microscopy of thin sections confirmed the presence of membrane septa separating individual compartments within filaments (Figure 2D), consistent with earlier reports in other filamentous Chloroflexota species (Gaisin et al., 2020). Interestingly, polar segments of filaments often appeared devoid of DNA, as evidenced by both fluorescence microscopy and electron microscopy (Figure 2E).

**Figure 2:**
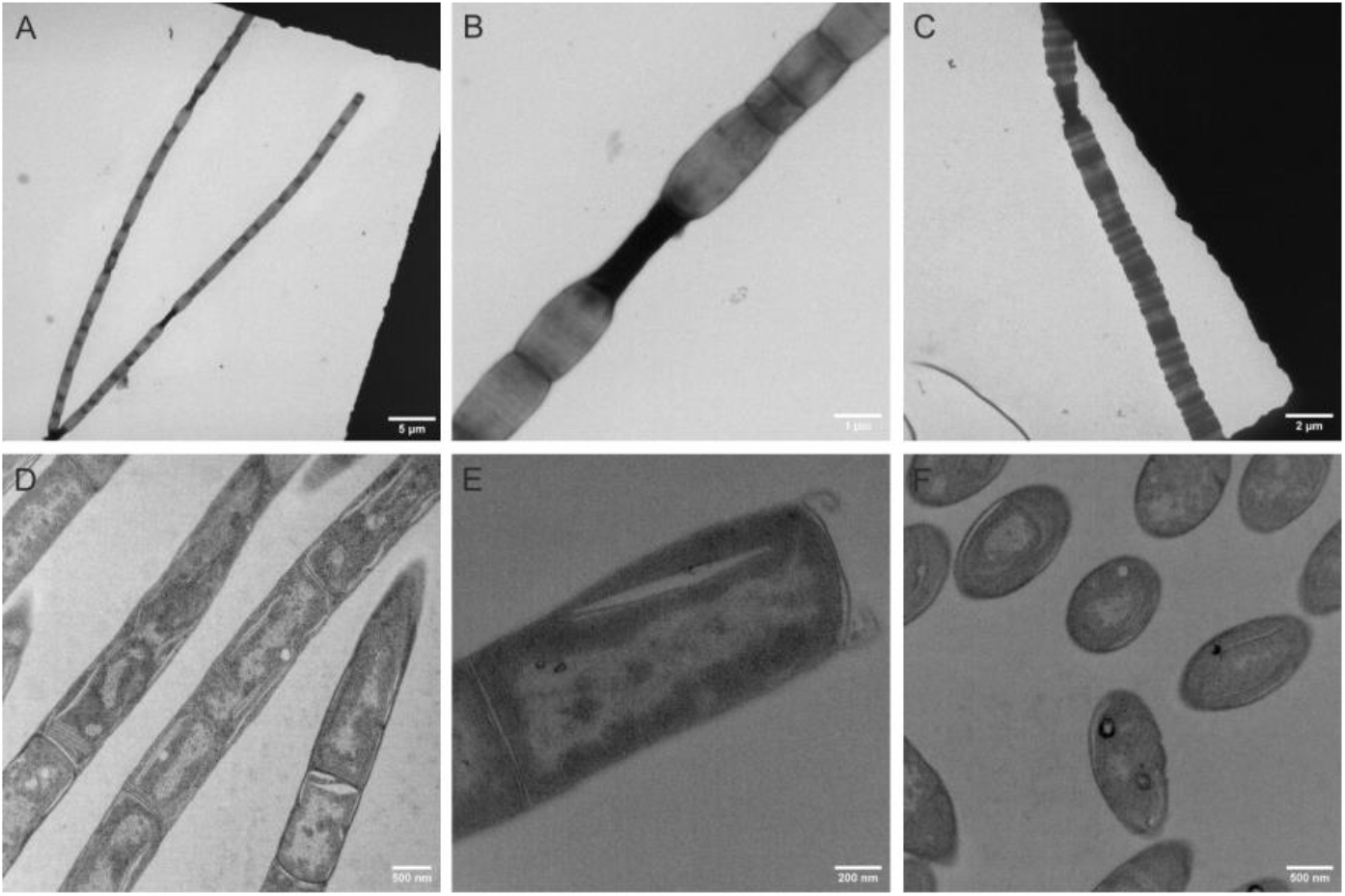
Electron microscopy images of *Litorilinea aerophila*. **(A-C)** Negative stained transmission electron microscopy images of *L. aerophila* captured in different growth phases: **(A)** lag phase, **(B)** exponential phase, and **(C)** stationary growth phase. **(D-F)** Ultra-thin sections of *L. aerophila* prepared for TEM. **(D)** Septal regions, **(E)** cell poles, and **(F)** lateral cross-sections of intracellular organization.

As cultures progressed through exponential and stationary phases, the overall filament length shortened. Membrane indentations became more regular, and smaller multicellular units were frequently detected (Figure 1B). Concurrently, cells developed round, surface-associated structures that stained positively with Nile Red, suggesting lipid-rich inclusions or vesicle formation. In the stationary phase (after 108 hours), filaments were further reduced in length, although no single cells were observed. Instead, vesicle-like membrane blebs were frequently seen budding from filament ends or indentation sites, indicating active membrane remodeling.

A particularly striking morphological phenotype was occasionally observed during stationary phase: filaments exhibiting a highly convoluted, tape-worm-like envelope morphology (Figure 1B, right panel). The biological function or structural basis of this phenotype remains unclear, but it was also detected in electron micrographs (Figure 2C), suggesting a reproducible and possibly stress-induced structural adaptation of the cell envelope.

### Cell division in *Litorilinea aerophila*

To date, live-cell imaging studies capturing filament growth and cell division dynamics have been reported for filamentous bacteria like *Streptomyces*, providing valuable insights into their spatial organization and division mechanisms (Jyothikumar *et al*., 2008; Schlimpert and Elliot, 2023). However, to our knowledge, no such observations have been made for members of the phylum Chloroflexota. Time-lapse imaging over 48 hours revealed that *L. aerophila* undergoes filamentous growth with asynchronous division events (Figure 3, Supplementary Figure 1, Supplementary Movie 2). A continuous flow of fresh medium ensured optimal, non-limiting growth conditions. The first constriction and subsequent division occurred after 4 hours, followed by the appearance of additional constriction sites at 8 hours. Interestingly, one of these sites produced a membrane bleb, suggesting potential vesicle formation or stress response. By 10 hours, division occurred at both sites, and continued elongation led to an additional division event by 14 hours (Figure 3, Supplementary Movie 1).

**Figure 3:**
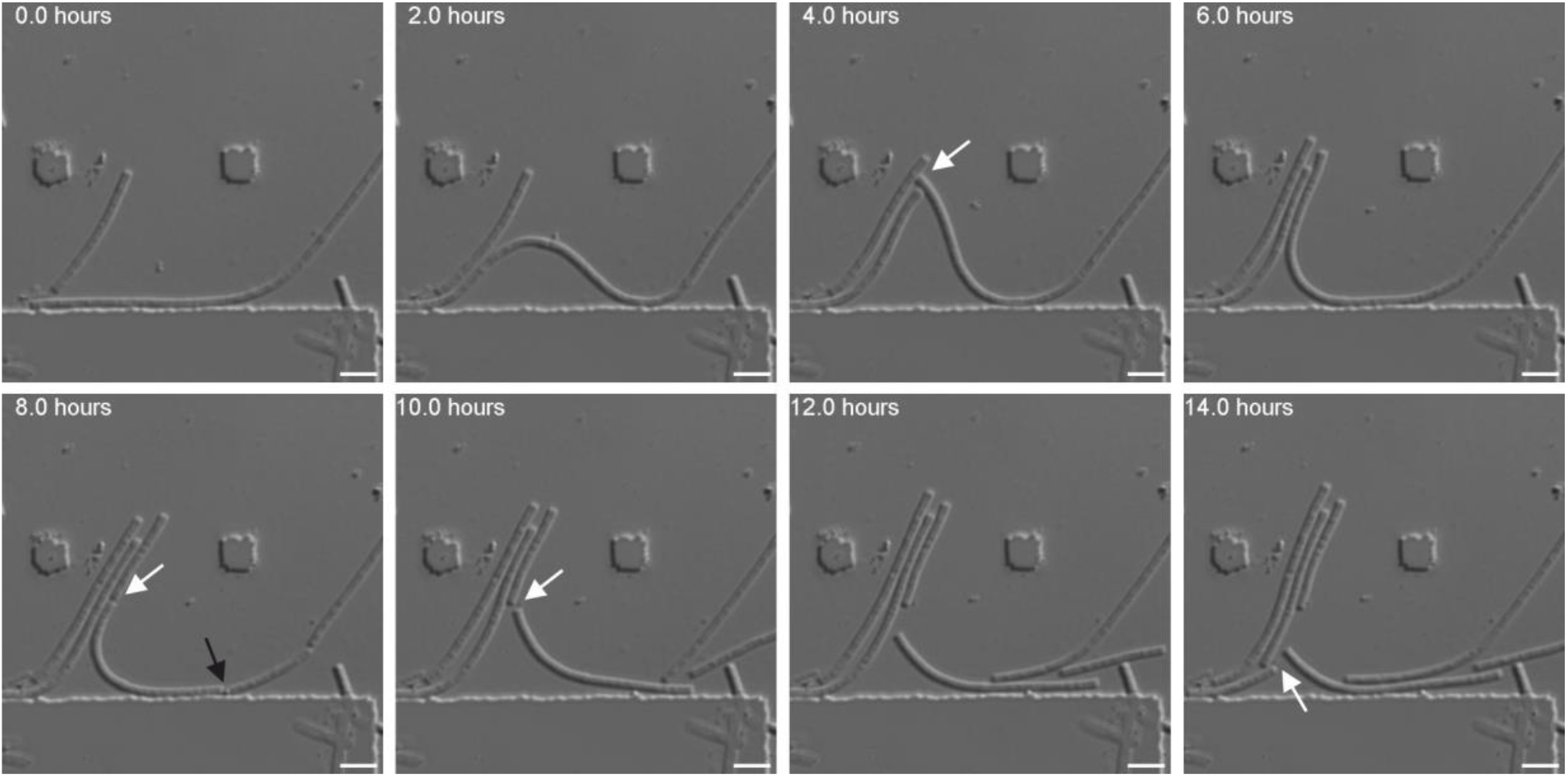
Cell division in *Litorilinea aerophila*. Live-cell microscopy of *L. aerophila* over a 14-hour period showing sequential division events (white arrows) within a filamentous structure. The first division occurs at 4 hours. Additional constriction sites appear by 8 hours, including one from which a membrane bleb is released (black arrow). At 10 hours, divisions occur at both constriction sites, followed by another division event observed at 14 hours. Scale bars: 10 µm.

### *Litorilinea aerophila* exhibits a multilayered cell envelope

To gain further insight into the ultrastructure of the cell envelope, cryo-electron tomography was done on vitrified samples of logarithmically growing *L. aerophila* cells. These cryo-electron tomograms revealed a multilayered cell envelope (Figure 4C-E). At the polar ends corresponding to the previously observed DNA-depleted regions, a distinct double-layered structure was observed (Figure 4C). Individual cells within the filament contained a well-defined internal structure. As the molecular composition of this structure remains unclear the term inner layer (IL) was chosen to refer to this structure. Each cell contained its own sacculus enclosed by this inner layer (IL, Figure 4D). Additionally, cryoET revealed the spherical, electron-dense structures within the inner layer that resemble storage granules as previously reported (Gaisin et al., 2020). Each cell was found to be enclosed by a cytoplasmic membrane (CM, Figure 4D). Within a filament, cells shared an additional distinct layer, which is described as the outer layer (OL, Figure 4D), as there is no clear evidence for a real outer membrane. The outer layer (OL) was further decorated by a continuous, proteinaceous layer (PL, Figure 4D) exhibiting characteristic lattice-like density that was detected surrounding the entire multicellular filament, consistent with the morphology of a surface (S)-layer. Adjacent cells were separated from each other by their respective inner layers within the filament (IL1 and IL2, Figure 4E). A connection between cell junctions between two cells was found reminiscent of septum channels found in other filamentous Chloroflexota (Gaisin *et al*., 2020). This septum channel (SC) connects the respective inner layers (IL) of the individual cells with each other (Figure 4E). In conclusion, we show a multilayered cell envelope architecture with an outermost layer encapsulating all cells within the filament, while individual cells have a separate layer. Early phylogenomic studies suggested a monoderm (single-membrane) organization, largely due to the absence of genes associated with outer membrane biogenesis (Sutcliffe, 2011). However, this assumption was challenged by Gaisin et al. (2020), who employed cryo-focused ion beam milling and cryoET to demonstrate a multilayered cell envelope in *Candidatus* Viridilinea mediisalina, *Chloroflexus aggregans*, and *Roseiflexus castenholzii* (Gaisin *et al*., 2020). These findings indicate that members of the phylum Chloroflexota may exhibit a broader range of envelope architectures than previously assumed.

**Figure 4:**
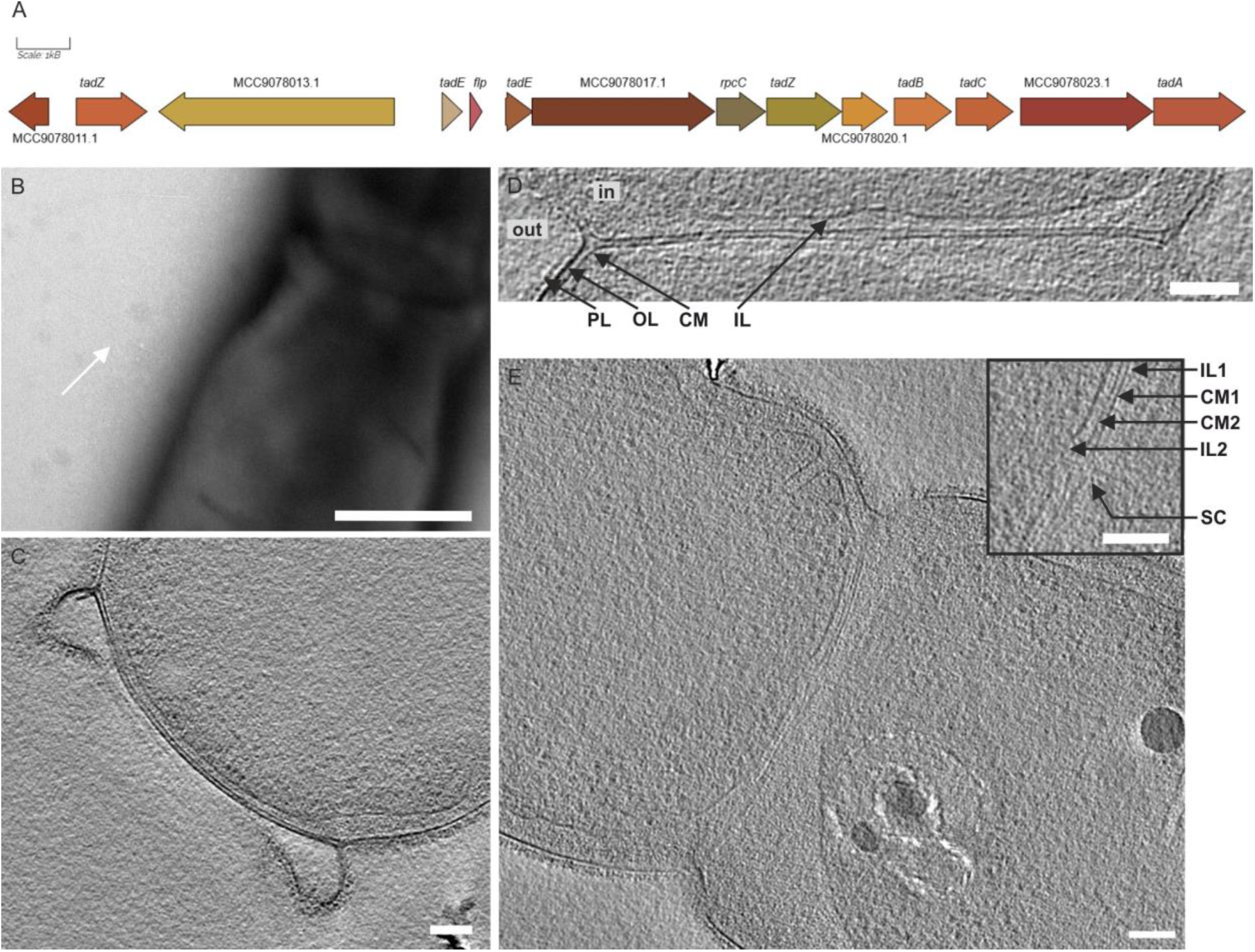
Cryo-Electron tomography. (A) Genetic organization of the *tad* gene cluster. (B) Negative-stained TEM images showing a Tad pilus (white arrow) on the lateral side of *L. aerophila*. (C) Cryo-electron tomogram of cell pole. (D) Close up of a cell-to-cell connection from a cryo-electron tomogram. (E) Cryo-electron tomogram from *L. aerophila* cells. The inset in the upper right corner shows a magnified view of the septum channel between two adjacent cells. Abbreviations: In, inside of the cell; Out, extracellular space; IL, inner layer; CM, cytoplasmic membrane; OL, outer layer; PL, protein layer; SC, septum channel. Scale bars: 500 µm (B), 100 µm (C-E).

### Identification of Tad Pili in Litorilinea aerophila

In addition to the previously described bacterial archaellum, a Type IV pilus homolog in *Litorilinea aerophila* (Sivabalasarma *et al*., 2025) negative-stain transmission electron microscopy revealed additional surface-associated filaments. Thin pilus-like structures were observed distributed along the lateral side of the cell (Figure 4B). Genomic analysis revealed the presence of a Tad-pilus operon including *tadA, tadB*, two copies of *tadZ, tadE, tadC* and one copy for *fpl* (Figure 4A, Supplementary Table 1). No additional pilus biogenesis systems were found, suggesting these cell appendages to be tad pili.

### A DUF11-containing protein forms grappling hook like structures

Additionally, to the tad pili, thicker filamentous structures were observed exclusively at the cell poles. These filamentous structures were approximately 180 nm in length and contained a grappling hook structure at their end. These resembled the grappling hook appendages recently described in *Aureispira sp*, which are formed by GhpA, a 5,898-residue monomer that spans the length of the filament and assembles as a heptamer (Lien *et al*., 2024). To identify a potential homolog in *L. aerophila* the GhpA sequence was used in a BLAST search against its genome, yielding a single hit: WP_141610714.1, which we will refer to as grappling hook like protein A (GhlA). This open reading frame encodes a DUF11 domain-containing protein with 6,596 amino acids and an estimated molecular weight of 672 kDa. Structural predictions using AlphaFold3 revealed a long filamentous structure composed of 49 DUF11 domains, arranged consecutively like beads on a string (Figure 5D). Notably, only 33 DUF11 domains are annotated on UniProt (ID: A0A540VE73_9CHLR). Each DUF11 domain consists of 10 antiparallel β-strands that assemble in two sheets, facing each other (Figure 5D). The structure closely resembles an immunoglobulin (IG)-like fold, which contains 7-9 antiparallel β-strands with two opposing sheets. The DUF11 domain-containing protein harbors a 19-residue hydrophobic stretch at the N-terminus that is predicted to be a transmembrane domain. Notably, it lacks an N-terminal signal peptide, implying that its secretion occurs via an alternative, signal peptide-independent pathway.

**Figure 5:**
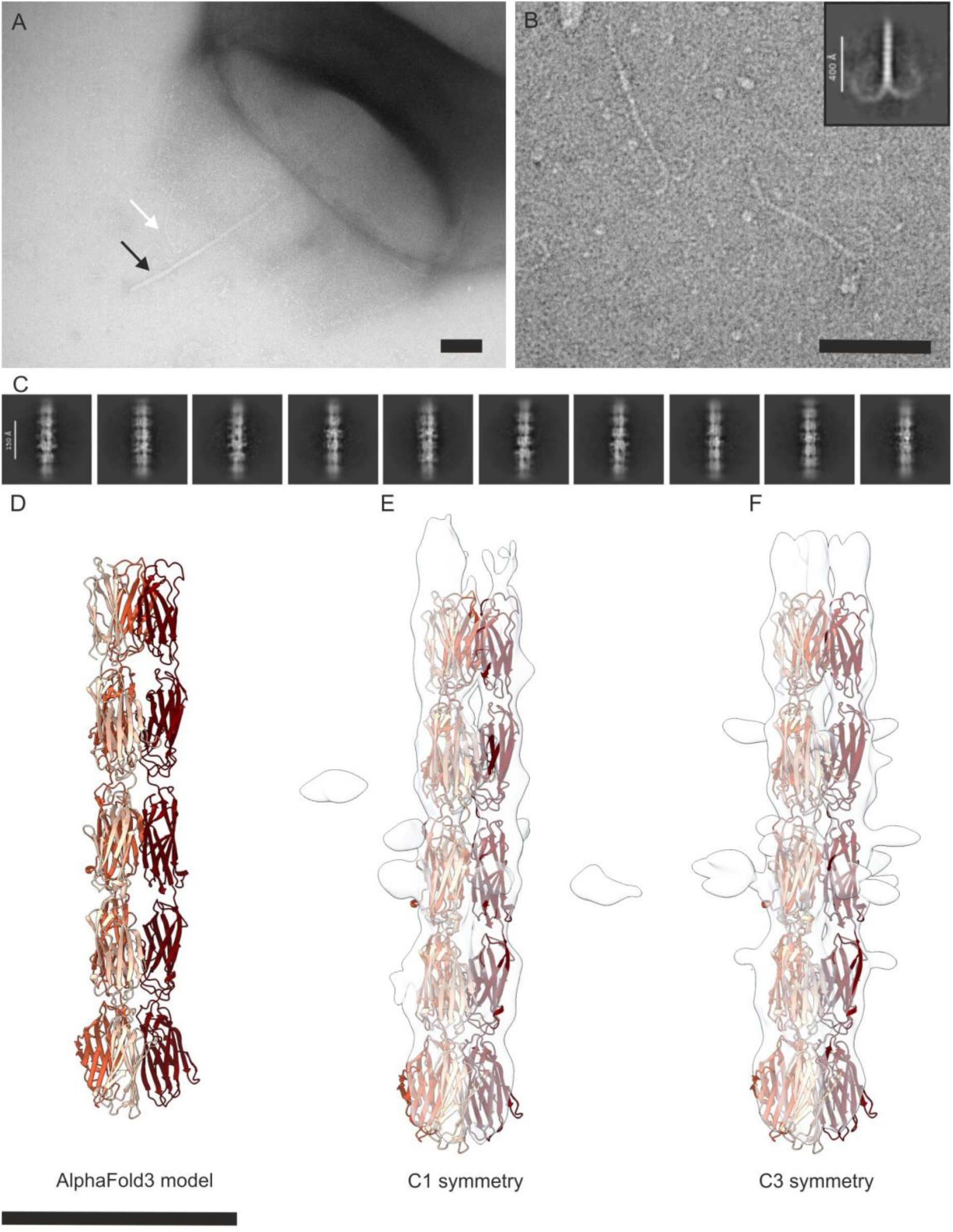
Grappling-hook structures. (A) Negative stain TEM image of cell poles of *L. aerophila*. Grappling hook-like structures (white arrow) and an archaellum (black arrow) are visible (B) Close-up of isolated grappling hook-like filaments imaged by TEM. An example 2D class average of negatively stained hooks is shown in the upper left corner. (C) 2D class averages of isolated hook filaments imaged by cryoEM. (D) Alphafold3 prediction of amino acid 2347-2954 of WP_141610714.1 corresponding to five DUF11 domains. The three different monomers are represented in different colors. (E) CryoEM volume from helical refinement job (white in surface representation), with no imposed symmetry (C1) and fitted AlphaFold3-model. (F) CryoEM volume from helical refinement job (white in surface representation), with imposed symmetry (C3) and fitted AlphaFold3-model. Scale Bars: 100 nm in panels A and B; 100 Å in panels D, E and F.

To verify the identity of the hook-forming protein, CsCl gradient fractions containing isolated hooks were analyzed by mass spectrometry. A total of 1272 proteins were identified (Supplementary Table 2). WP_141610714.1 was identified with 26 unique peptides located between amino acids 257 and 6566 of its predicted sequence (Supplementary Table 3) and ranked 38th in overall intensity (Supplementary Table 2). Since large proteins tend to be overrepresented in peptide-centric quantitative MS, intensity-based absolute quantification (iBAQ) values were calculated for all proteins from summed peptide intensities. This allows for the estimation of relative copy numbers and, taking into account theoretical molecular weights, relative weight fractions for all proteins. WP_141610714.1 was ranked 182nd in estimated relative molecule numbers and 21st in weight fraction among all identified proteins (Supplementary Figure 2, Supplementary Table 3).

### Structural analysis of the grappling hook Filament

For further structural analysis, hook filaments were vitrified on cryoEM grids and imaged, resulting in a set of 11.060 movies. Motion correction and patch-based CTF estimation were carried out in CryoSPARC. Particle picking was performed using a CrYOLO model trained on 40 manually annotated micrographs, yielding 1.858.183 particles. Due to the high flexibility of the tip of the filamentous hook structure, we decided to focus on the core of the hook and neglected the grappling hook region (Figure 5B). Particles were extracted from motion-corrected, CTF-estimated micrographs and Fourier cropped from 100 to 300 pixels. After four rounds of 2D-classification, 78 classes were obtained, including 519,123 particles. Most of the classes were blurry, likely due to separate filament regions getting averaged into one class. However, 10 classes, including 104,631 particles, contained sharp, well-distinguishable features. The best 2D class averages achieved resolutions between 7 Å and 9 Å (Figure 5C). The DUF11 domains were distinctly resolved and, consistent with the structure prediction, appeared as beads on a string-like structures. Some of the single DUF11 domains were tilted, relative to other DUF11 domains. It remains unclear whether these variations represent fixed features or whether they indicate some conformational flexibility between the domains. Additionally, the DUF11 domains appeared O-shaped, which may result from a gap between the opposing β-sheets within the IG-like fold. Furthermore, densities protruding from the filament were observed, that could not be assigned to the previously AlphaFold3-predicted structure. As these densities are too large to represent glycan modifications, they could either be proteinaceous features or simply amplified noise.

Using the sharpest 10 classes, an ab-initio job was run for initial 3D-reconstruction. The output volume was used as an initial input for another helix refine job, which was run with C1 (unbiased) symmetry, producing a volume with a resolution of 6.75 Å (Figure 5E). The volume strongly suggested three strands; therefore, a helix refine job with a biased C3 symmetry was run and produced another volume, improving the resolution to 6.39 Å (Figure 5F). Other symmetries (C2-C10) were experimentally imposed but did not lead to structures of convincing quality. To further support the assignment of C3 symmetry, an AlphaFold3-model was generated, using three copies of the residues 2347-2954 (corresponding to 5 DUF11 domains) WP_141610714.1 (Figure 5D). The resulting model could be fitted into both C1 and C3 volumes (Figure 5E,F). This indicated a probable C3 symmetry of the hook-like filament.

## Discussion

*Litorilinea aerophila* shows multicellular organized filaments, in which individual cells are separated by membrane indentations (Figure 1,2), consistent with the initial description by (Kale *et al*., 2013). Frequently, membrane blebs were observed in light microscopy. Combined with the observation of DNA-depleted polar regions within the filaments, these membrane blebs are interpreted as membranes shed from filament ends, potentially representing membrane fragments left behind following cell rupture.

Filamentous cyanobacteria are known to differentiate into specialized cell types along the filament, often exhibiting distinct morphologies (Risser, 2023). However, such differentiation was not evident in *L. aerophila* using light microscopy or thin-section electron microscopy, suggesting either the absence of morphologically distinct cells or limitations of the applied imaging methods. Notably, during growth *L. aerophila* filaments transitioned from long, irregular segmented structures into shorter filaments with more uniformly spaced membrane septa. A similar diversity of filament length has been reported in Cyanobacteria, though the regulation of the filament length is not clearly understood. It has been suggested that longer filaments contribute to biofilm or microbial mat formation under favorable conditions, while shorter filaments are released as an environmental stress response (Risser, 2023). This behavior is exemplified by the formation of hormogonia, 5-25 cells short and motile filaments in species such as *Anabaena, Nostic, Oscillatoria* and *Trichodesmium spp*. (Seki *et al*., 1981; Smith and Gilbert, 1995; Kruskopf and Plessis, 2006; Silva *et al*., 2025). In *Anabaena*, hormogonium release is triggered by nutrient limitation and temperature shifts, a mechanism that may also be relevant for *L. aerophila* (Smith and Gilbert, 1995). In liquid cultures, longer *L. aerophila* filaments formed dense cell aggregates resembling early-stage biofilm structures. As growth continued, these filaments shortened and, as reported previously, became motile (Sivabalasarma *et al*., 2025). Thus, this morphological transition could facilitate the search for environmental-friendly habitats under nutrient-limiting conditions, similar to the hormogonium-based strategy in cyanobacteria. Unlike filamentous cyanobacteria, where filament elongation happens apical by the addition of a single cell, *L. aerophila* did not show evidence of tip-based growth (Silva *et al*., 2025). Time-lapse light microscopy revealed elongation occurring along the entire length of the filament rather than at the apical end (Supplementary Movie 1). This mechanism is known from cable bacteria, which also display non-apical filament elongation, possibly due to the high growth rates incompatible with tip growth (Schauer *et al*., 2014). Genomic analysis of *L. aerophila* reveals an FtsZ-based cell division system. In cable bacteria, cell division occurs synchronously across entire filaments, which can exceed 700 cells in length (Geerlings *et al*., 2021). Synchronous division has also been described in *Anabaena* (Corrales-Guerrero *et al*., 2018). In contrast, *L. aerophila* exhibited asynchronous division events that appeared as irregular or asymmetric cellular breakages (Figure 3). This variability highlights the broader diversity of division strategies among filamentous prokaryotes. For example, *Trichodesmium erythraeum* and *Oscillatoria* spp. do not undergo synchronized division (Popa *et al*., 2007; Sandh *et al*., 2009). Similarly, asymmetric division has been documented in filamentous forms of *Synechococcus elongatus* under dim light conditions; however, these structures are elongated single cells rather than true multicellular filaments (Liao and Rust, 2018)

The cell envelope architecture in Chloroflexota has historically been considered monoderm, largely due to the absence of canonical outer membrane markers such as the BAM complex and lipopolysaccharide biosynthesis genes (Sutcliffe, 2010). However, our findings on *L. aerophila* challenge this classification and contribute to the growing evidence that Chloroflexota exhibit a broader diversity in envelope organization than previously thought. Cryo-electron tomography analysis reveals a complex, multilayered envelope structure in *L. aerophila*, composed of an inner layer, a cytoplasmic membrane, an intermediate compartment bounded by an outer layer, and a rough external surface (Figure 4D). This architecture is highly reminiscent of that observed in *R. castenholzii*, a member of the Chloroflexia class, despite the phylogenetic distance between the two organisms. The similarities include the absence of a distinct intermediate layer as seen in *Ca*. Viridilinea mediisalina and *C. aggregans*, as well as the direct connection of the septum to the outer layer (Gaisin *et al*., 2020). Although our data do not resolve the molecular composition of the outer and external layers, the structural resemblance to peptidoglycan-rich layers in *R. castenholzii* and the presence of thick septa in *L. aerophila* suggest a potentially modified or specialized cell wall organization. It is particularly noteworthy that neither organism encodes known outer membrane biosynthetic genes, implying that the observed layers may represent convergent evolutionary adaptations rather than homologous diderm envelopes. Interestingly, while *Ocs. trichoides* contains peptidoglycan with ornithine instead of the typical diaminopimelic acid, no such chemical data are currently available for *L. aerophila* (Keppen *et al*., 2018). The presence or absence of similar atypical PG structures in *L. aerophila* remains to be determined and would offer valuable insights into the evolution of envelope complexity in this phylum.

*In addition to the archaellum described previously in L. aerophila, Tad pili were observed at the cell poles. Several genes associated with the tad pilus biogenesis system were identified in L. aerophila* (Figure 4). Tad pili are known to mediate twitching motility, DNA uptake, surface adhesion, and biofilm formation, and likely contribute to the observed aggregation behavior of *L. aerophila in liquid culture*. Phylogenetic studies suggest that the Tad system originated from archaeal Epd-type type IV pili and was laterally transferred to bacteria (Denise *et al*., 2019). The genetic similarities between Tad and archaeal pili systems—including homologous membrane components and evidence of gene fission—support an archaeal origin for the Tad machinery in *L. aerophila*. Additionally, hook-like cell surface structures were identified (Figure 5A,B), resembling the predatory grappling hooks described in *Aureispira sp*, where they are implicated in ixotrophic behavior (Lien *et al*., 2024). To understand the function of the hook filaments in *L. aerophila*, the genome was searched for other related hook- and ixotrophy genes. Homologs of the outer membrane basal body of the *Aureispira sp*. hook *ghpB* and *ghpC* were not found in *L. aerophila*, which is in accordance with the missing secondary membrane. The genomic neighborhood of the *L. aerophila* grappling hook like protein *ghlA* was inspected for similar genes but showed no interesting hits. Apart from bacteria, similar hook-like structures have also been found in archaea with quite a different functionality. These structures, called hami, are known to facilitate the formation of interconnected networks between cells (Moissl *et al*., 2005; Probst *et al*., 2014). Even though visually hami tips in particular look very similar to *L. aerophila* hooks, no homologs of the hamus subunit protein (Moissl *et al*., 2005; Perras *et al*., 2015) could be found in *L. aerophila*. Therefore, these structures likely arose independently through convergent evolution.

Different imaging approaches revealed a combination of non-apical filament elongation, asynchronous cell division and multilayered cell envelope in *Litorilinea aerophila*. This challenges the conventional concepts of bacterial multicellularity and cell envelope organization. The presence of Tad pili and hook-like surface structures points towards functional adaptations related to motility, adhesion and intercellular interaction. Altogether, these findings expand the current understanding of cell envelope diversity in Chloroflexota.

^19,61,63,69,70^, therefore, *L. aerophila* cells were spotted on semi-solid agar plates. After five days, a thin halo appeared and cells from the rim and midpoint of the halo were imaged with light and fluorescence microscopy (Figure 4). Cells from motility plates were mostly shorter (4-10 µm), yet remained multicellular, as shown by fluorescence microscopy (Figure 4). Individual cells of a multicellular filament were separated through a septum (Figure 4a). Electron microscopy imaging of the short filaments of cells revealed surface appendages with 11-12 nm in diameter, exclusively found at the cell poles of shorter multicellular filaments (Figure 4b). Surface appendages were present on cells isolated from the rim of the motility plate as well as from the midpoint (Figure 4b). Additionally, qRT-PCR showed the expression of the archaellin encoding gene *arlB* as well as the archaellum machinery genes in cells from semi-solid agar plates compared to liquid medium grown cells (Supplementary Figure 3).

Given the presence of the archaellum filament, the swimming motility of *L. aerophila* was imaged by time-lapse light microscopy. Swimming cells were analyzed and tracked using TrackMate7^35^. Cells from a motility plate displayed motile behavior with a swimming speed of 10.46 ±6.68 µm/s (Supplementary Video 1). This presents the first actively swimming member of the Chloroflexota phylum and, notably, the first example of a bacterium utilizing an archaellum for motility.

## Methods

### Bacterial strains

*Litorilinea aerophila* PRI 4131 (DSM 25763) was obtained from the DSMZ (German Collection of Microorganisms and Cell Cultures GmBH). *Litorilinea aerophila* PRI 4131 was previously isolated from an intertidal hot spring in Isafjardardjup, NW Iceland (Kale *et al*., 2013).

### Growth evaluation

*Litorilinea aerophila* was grown in 5 ml Marine Broth (MB) (Difco 2216) in plastic tubes. Cells were incubated at 55°C with shaking at 90 rpm. For determination of the cell dry weight (mg ml-1) one 5 ml culture was inoculated for each measuring point to a theoretical OD 0.02. Cell were grown at 55°C, 90 rpm shaking. 4 ml of these cultures were pelleted, and the cells were washed once with PBS. The cell pellet was transferred to calibrated reaction tubes and dried in a speed vac at 45°C for one hour before measuring the weight of each pellet. The experiment was done in triplicate.

### Light and Fluorescence Microscopy

0.5 µl Syto13 (ThermoFisher) and 1 µl Nile red (5 mg/ml in DMSO) (ThermoFisher) were added to 500 µl of each culture from the dry weight determination. Cells were incubated for 10 min in the dark at room temperature. Cells were washed once with PBS, followed by resuspension in 500 µl PBS. For imaging 5 µl of cells of spotted on an agarose pad made of 1% agar in PBS. Cells were observed at x100 magnification in the DIC mode using an inverted microscope (Zeiss Axio Observer.Z1, controlled via Zeiss Blue v.3.3.89 software). Image analysis was performed using ImageJ.

### Negative stain electron microscopy

5 ml Marine Broth medium was inoculated with *L. aerophila* and grown overnight at 55°C, 90 rpm shaking. Samples were taken from day 1,2,3 and 5 µl of cells were applied on freshly glow-discharged carbon/Formvar-coated copper grids (300 mesh; Plano GmbH) and incubated for 30 s. The excess liquid was blotted away and the grid washed three times with ddH2O, followed by negative staining with 2% uranyl acetate. Images were taken with a Hitachi HT7800 operated at 100 kV, equipped with an EMSIS Xarosa 20-megapixel CMOS camera.

### Thin-section preparation for electron microscopy

5 ml of cell culture were centrifuged for 5 min at 5.000 rpm. The cell pellet was resuspended in 50 µl of 2% (w/v) Ultra-low Gelling Temperature Agarose (SIGMA A-2576) and let solidify at 4°C for 5 min. Cells embedded in the agarose were fixed in a 2,5 % (v/v) glutaraldehyde solution in MTSB Buffer (50 mM PIPES, 5 mM EGTA, 5 mM MgSO_4_·7H_2_O, 50 mM KOH) for 5 h at room temperature and overnight at 4°C with fresh fixative solution. Samples were washed with MTSB five times (10 min each) and post fixed in aqueous 1% (w/v) OsO_4_ on ice during 3 h. Samples were washed at room temperature with water five times, each for 10 min, and *in bloc* stained with 2% UrAc (w/v) in water for 2 h. After washing the samples twice with water for 5 min, they were dehydrated by incubation for 15 min in increasing EtOH/water graded series (30%. 50%, 70%, 80%, 95%) and for 30 min twice in 100% EtOH and 100% Acetone. Samples were gradually embedded in resin by incubating them in Agar 100:acetone mixtures (1:3, 1:1, 3:1) 8 h each, and finally inpure resin (3 times exchange, 8 h each). Resin blocks containing the embedded cells were polymerized for two days at 60°C and 70 nm sections were obtained with a Reichert-Jung Ultracut-E microtome. Sections were collected on copper grids, contrasted with 2% uranyl acetate and Reynolds lead citrate solution(Reynolds, 1963) and observed in a Hitachi 7800 TEM coupled to a Xarosa CMOS camera (Emsis).

### Purification of cell surface filaments

A preculture was grown by inoculating a single colony of *Litorilinea aerophila* into 5 mL MB and incubating overnight at 55 °C with gentle shaking (90 rpm). The following day, 1 mL of culture was transferred into 2 × 50 mL MB in 250 mL Erlenmeyer flasks and grown overnight under the same conditions. The OD_600_was measured and diluted to ~0.01–0.02 into 4–6 × 400 mL MB in 2 L flasks, followed by incubation at 55 °C for 5 days with 90 rpm shaking. Cells were harvested by centrifugation (3000 x g, 30 min), resuspended in 50 mL isolation buffer (1x PBS +2% NaCl), and subjected to hook shearing using a peristaltic pump: 1 h through a 0.9 mm × 40 mm needle, then 2 h through a 0.45 mm × 10 mm needle, both at 90 rpm. Sheared cells were removed by low-speed centrifugation (12,000 x g, 25 min, 4 °C), and the supernatant was ultracentrifuged (200,000 x g, 1 h 20 min, 4 °C). The resulting pellet was resuspended in 500 µL isolation buffer and layered onto a 4.5 mL CsCl gradient (0.5 g/mL in isolation buffer), followed by overnight centrifugation (250,000 x g, 16 h, 4 °C). 500 µL fractions were collected, diluted to 8 mL with isolation buffer, and pelleted again (250,000 x g, 1.5 h, 4 °C). Pellets were resuspended in 50 µL isolation buffer and analysed by negative-stain TEM to identify hook-rich fractions. Selected fractions were concentrated to 25 µL using 100 kDa Amicon filters (Sigma Aldrich).

### Cryo-electron microscopy sample preparation and data acquisition

R2/2 copper grids (Quantifoil Micro Tools) were glow discharged and 3.5 μL of isolated hook filaments were applied. The grids were vitrified in liquid ethane and stored in liquid nitrogen. The dataset was collected with a Titan Krios electron microscope (Thermo Fisher), equipped with a Selectris energy filter (Gatan) and a Falcon 4i electron detector (Thermo Fisher). For data collection the pixel size was calibrated to 1.22 Å, corresponding to a magnification of 105,000 x. Using EPU 3.6 software (Thermo Fisher), 11,060 movies were collected as 40-fraction movies with an exposure time of 5.85 s, a total electron dose of 40 e-/ Å 2 and a defocus range of −0.5 to −2.2 µm.

### Processing of the hook filaments dataset in CryoSPARC

Image processing was performed in CryoSPARC v4.6 (Punjani *et al*., 2017). Motion correction and CTF estimation were done using the patch-based methods. An initial set of 30 manually picked micrographs was used to train a CrYOLO filament-picking model in filament mode (Wagner *et al*., 2019, 2020), resulting in 1,858,183 automatically picked particles. Particles were extracted with a 300 px box size and Fourier-cropped to 100 px for four rounds of 2D classification (40 iterations, 100 classes, batch size 500). From this, 519,123 particles across 78 classes were selected and re-extracted at full resolution. An initial 3D model was generated ab initio from the sharpest 104,631 particles (10 classes) using C1 symmetry and windowing (0.75–0.9), with minibatch sizes of 300–1000. This was refined using helical refinement first with C1 (6.75 Å), then C3 symmetry, supported by fitting an AlphaFold-predicted trimer (WP_141610714.1) (Jumper *et al*., 2021). The full 519,123-particle set was then refined using this C3 volume, improving the resolution to 5.05 Å. To explore heterogeneity, 3D classification (five classes, 7 Å filter, C3 symmetry, hard classification) was followed by helical refinement of each class (5.97–6.23 Å, ~86k– 128k particles). The best class was further classified and refined, but yielded lower-resolution volumes (7.1–7.53 Å, ~17k–26k particles). A strand-specific mask was generated in ChimeraX (Goddard *et al*., 2018) and applied in a helical refinement (C1, 519,123 particles), yielding only minor resolution gains. Attempts to loosen the mask via threshold/dilation worsened the results.

### Mass spectrometry sample preparation, data acquisition and analysis

Proteins were denatured in 1% SDS and cysteine residues were reduced with 10 mM DTT for 15 min at 95°C. For alkylation of free thiol groups 50 mM chloroacetamide was added for 30 min at 22°C. The reaction was quenched with 50 mM DTT. Samples were purified and digested with trypsin in a 96-deep-well plate (1 mL, Agilent Technologies) with an SP3-bead-based protocol (Hughes *et al*., 2019) adapted to a STARlet automated liquid handling system (Hamilton, Bonaduz Switzerland) equipped with a 96-well magnet plate (Magnum Flex, Alpaqua). Proteins were bound to 100 µg Sera-Mag SpeedBead magnetic carboxylate-modified particles (Cytiva) with 80% ethanol for 20 minutes at room temperature and washed three times with 80% ethanol. Proteins were digested overnight at 37°C in 50 µL of 100 mM ammonium bicarbonate containing 0.15 µg of trypsin (Promega V5111, Madison, USA). Digestion was stopped by adding 30 µL of 5% formic acid (FA). Peptides were desalted using self-packed SDB-RPS Stop and Go Extraction tips (StageTips) (Rappsilber *et al*., 2007) composed of three layers of 1.0 × 1.0 mm (AttractSPE® Disks Bio RPS, Affinisep). The StageTips were equilibrated sequentially with 100% methanol, 80% acetonitrile containing 0.1% FA, and twice with 0.1% FA. Acidified peptide solutions were loaded onto the StageTips and washed with 0.1% FA, 80% acetonitrile with 0.1% FA, and 80% methanol with 0.5% FA. Peptides were eluted using 60% acetonitrile containing 5% ammonia, dried in a vacuum centrifuge and reconstituted in 0.1% TFA.

LC-MS analysis was carried out on an UltiMate 3000 RSLCnano system coupled to a Q Exactive mass spectrometer (both Thermo Fisher Scientific) as described (Bhuiyan *et al*., 2025)with minor changes. In brief, peptides were preconcentrated on a PepMap C18 trapping column and separated on a μPAC™ C18 pillar array column (200 cm bed length, Thermo Fisher Scientific) applying a binary buffer system consisting of A (0.1% formic acid) and B (86% acetonitrile, 0.1% formic acid) and a linear gradient from 5% to 50% B in 45 min and 50 to 99% B in 2 min at a flow rate of 1 µl/min. Cycles of data-independent acquisition (DIA) consisted of: one overview spectrum (RF lens of 50%, normalized AGC target of 3e6, maximum injection time of 60 ms, *m/z* range of 385 to 1.050, resolution of 70.000 at *m/z* 200, profile mode) followed by MS2 fragment spectra (RF lens of 50%, normalized AGC target of 1e6, maximum injection time of 60 ms) generated sequentially by higher-energy collision-induced dissociation (HCD) at a normalized energy of 26% in a precursor *m/z* range from 400 to 1.000 in 50 windows of 12.5 *m/z* isolation width overlapping by 0.25 *m/z* and recorded at a resolution of 35.000 in profile mode.

Mass spectrometric raw data were converted to mzML format using the ProteoWizard software, version 3.0.21229 (Chambers *et al*., 2012). For protein and peptide identification a library free search was performed using FragPipe, version 22.0 (Yu *et al*., 2023) with quantification by Dia-NN, version 1.82 (Demichev *et al*., 2020) against the organism specific sequence database for *Litorilinea aerophila* from Uniprot database (release 2025_02, 4877 entries). Trypsin was set as protease with 2 allowed missed cleavages. Carbamidomethylation of cysteine was selected as fixed modification, N-terminal excision of methionine, oxidation of methionine and N-terminal acetylation were selected as variable modifications with a maximum number of 3. Further parameters were a 1% protein false discovery rate cutoff, unrelated runs and robust LC quantification. All protein groups with non-zero intensities listed in the protein and protein group output matrices of DIA-NN were selected for further analysis. Intensity-based absolute quantification (iBAQ) values were calculated by dividing the sum of peptide intensities by the theoretical number of peptides predicted for proteolytic digestion by trypsin, considering peptides in the range of 7 to 35 amino acids that typically detected during LC-MS analysis (Schwanhäusser *et al*., 2011). Relative weight fractions were derived by multiplying iBAQ values for each protein with its predicted molecular weight and normalizing to the sum across all proteins.

### Cryo-electron tomography

*L. aerophila* culture was grown to OD_600_~0.9 in liquid marine broth medium at 55°C and 90 rpm. A total of 3.5 µl cells were applied on freshly glow-discharged EM copper grids (R2/2, Quantifoil) and plunge frozen in liquid ethane using the Leica GP2. Frozen grids were stored in liquid nitrogen. CryoET data was collected using a Titan Krios (Thermo Fisher Scientific) cryo electron microscope operating at 300 kV equipped with a Falcon4i and Selectris energy filter (Gatan) at a calibrated pixel size of 4.71Å corresponding to 26 000x magnification. Using Tomography 5 (Thermo Fisher Scientific) 12 tilt series were collected with a tilt range of +60° to −60° in 3° increments, a total electron dose of 140 e_-_/ Å_2_ and a defocus range from −5 - −8 µm. Tilt series were driftcorrected using alignframes in IMOD (Kremer *et al*., 1996). Corrected tilt series were used to reconstruct 4x binned cryo-tomograms with IMOD using weighted back projection. Tomograms were CTF-deconvolved and filtered using isonet (Liu *et al*., 2022).

## Data availability

The mass spectrometry proteomics data have been deposited to the ProteomeXchange Consortium via the PRIDE partner repository (Perez-Riverol *et al*., 2025)with the dataset identifier PXD066799 (Reviewer access details: Project accession: PXD066799; Token: mz0A1Tb4XsDJ). The cryoEM map have been deposited in the EMDB under accession numbers EMDB-54772 (C1 symmetry) and EMDB-54773 (C3 symmetry). The cryo-electron tomograms have been deposited in the EMDB under accession number EMDB-54781 (cell pole) and EMDB-5780 (cell).

## Acknowledgments

We want to thank Rosula Hinnenberg of the EM facility at the Faculty of Biology, University of Freiburg, for her assistance during the generation of EM data. The TEM (Hitachi HT7800) was funded by the DFG grant (project number 426849454) and is operated by the University of Freiburg, Faculty of Biology, as a partner unit within the Microscopy and Image Analysis Platform (MIAP) and the Life Imaging Center (LIC), Freiburg. We would also like to thank Stefan Steimle at the Freiburg CryoEM facility for help with cryoEM screening and data collection. Electron microscopy data were collected at the Cryo-EM Facility of the University of Freiburg. The Titan Krios G4 cryoTEM used for imaging was funded by Deutsche Forschungsgemeinschaft (project no. 506518771) and is operated within the Microscopy and Image Analysis Platform (MIAP), University of Freiburg. We thank Carola Hunte for providing access to their plunge freezer. We want to thank Martin Pilhofer and Davide Amendola from the Pilhofer Lab for helpful discussions regarding data processing.

