## Supplementary material for "Filamentous growth, cell envelope architecture, and surface appendages of a member of the Chloroflexota, *Litorilinea aerophila*"

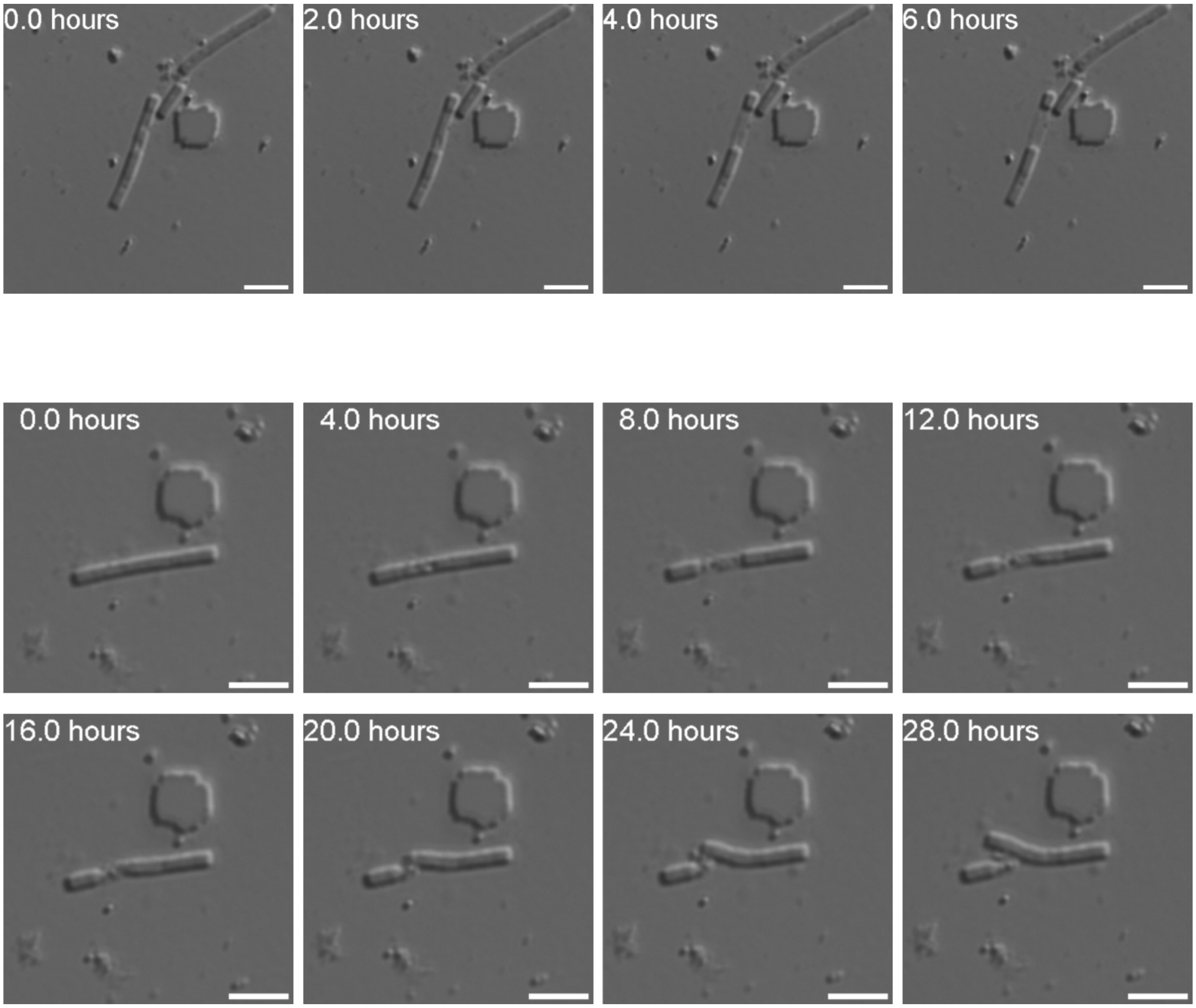


**Supplementary Figure 1. Time-lapse microscopy of *L. aerophila* cell division.** Two representative time-lapse series show sequential stages of division, highlighting morphological changes from elongation to septum formation and daughter cell separation. Time stamps indicate hours post-imaging initiation.

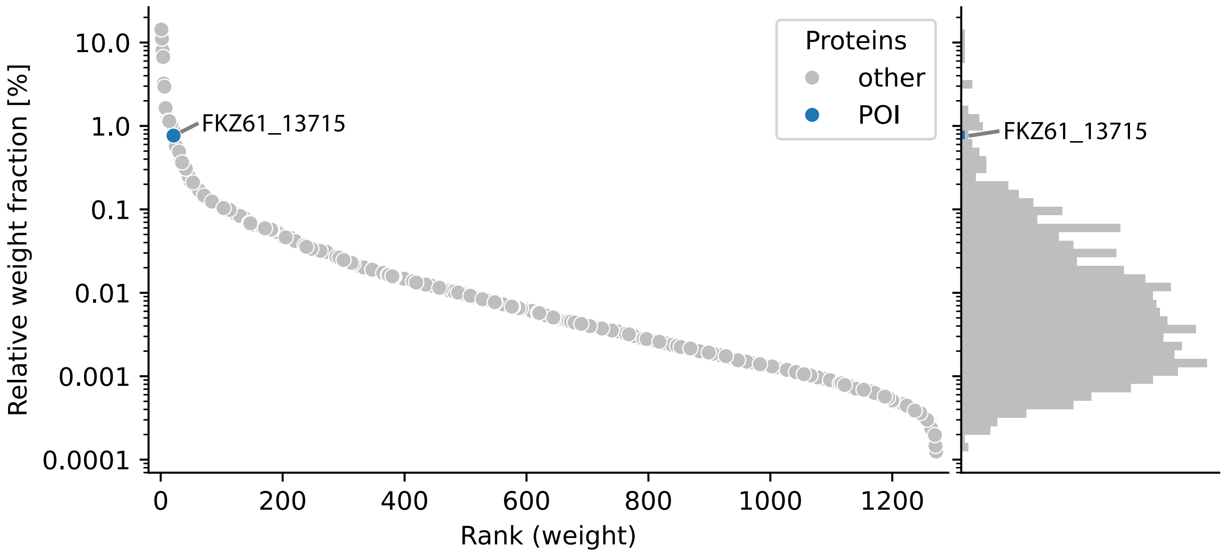


**Supplementary Figure 2.** Quantitative MS analysis of CsCl gradient fractions of sheared filaments from L. aerophila. Ranking of all proteins by their relative weight fraction based on intensity-based absolute quantification (iBAQ) places WP_141610714.1 as 21$^{st}$.

**Supplementary Table 1.** Tad-Pilus encoding genes from the *L. aerophila* genome. The table contains an overview of the found proteins, accession numbers, residue count, an annotated description and possible function.

| Protein | Accession Nr. | Residues | Annotated Description | Possible Function |
|---|---|---|---|---|
| **Flp/PilA** | MCC9078015.1 | 66 | Flp family type IVb pilin | Major pilin |
| **TadE/CpaJ 1** | MCC9078014.1 | 110 | Pilus assembly protein | Minor pilin |
| **TadE/CpaJ 2** | MCC9078016.1 | 148 | Pilus assembly protein | Minor pilin |
| **TadZ/CpaE 1** | MCC9078012.1 | 386 | Response regulator | ParA/MinD ATPase |
| **TadZ/CpaE 2** | MCC9078019.1 | 407 | AAA family ATPase | ParA/MinD ATPase |
| **RcpC/CpaB** | MCC9078018.1 | 267 | Flp pilus assembly protein CpaB | Inner membrane subunit |
| **TadB/CpaG** | MCC9078021.1 | 311 | Type II secretion system F family protein | Platform protein |
| **TadC/CpaH** | MCC9078022.1 | 310 | Type II secretion system F family protein | Platform protein |
| **TadA/CpaF** | MCC9078024.1 | 499 | CpaF family protein | Motor ATPase |
